# Dopamine and stress signalling interplay patterns social organization in mice

**DOI:** 10.1101/856781

**Authors:** Dorian Battivelli, Cécile Vernochet, Claire Nguyen, Abdallah Zayed, Aura Carole Meirsman, Sarah Messaoudene, Alexandre Fieggen, Gautier Dreux, Fabio Marti, Thomas Contesse, Jacques Barik, Jean-Pol Tassin, Philippe Faure, Sébastien Parnaudeau, François Tronche

## Abstract

The rules leading to the emergence of a social organization and the role of social hierarchy on normal and pathological behaviours remain elusive. Here we show that groups of four isogenic male mice rapidly form enduring social ranks in a dominance hierarchy. Highest ranked individuals display enhanced anxiety and working memory, are more social and more susceptible to stress-related maladaptive behaviours. Are these differences causes or consequences to social life? We show that anxiety emerges from life in colony whereas sociability is a pre-existing trait. Strikingly, highest ranked individuals exhibit lower bursting activity of VTA dopamine neurons. Both pharmacogenetic inhibition of this neuronal population and the genetic inactivation of glucocorticoid receptor signalling in dopamine-sensing brain areas promote the accession to higher social ranks. Altogether, these results indicate that the shaping of social fate relies upon the interplay of dopamine system and stress response, impacting individual behaviour and potentially mental health.

## Introduction

Social organization is readily observable across vertebrate species and can result in the establishment of a social hierarchy that may minimize energy costs due to direct competitions for resources among congeners (Tinbergen, 1939; Francis, 1984). At the group level this may improve adaptation to the environmental demands. At the individual level, it exposes different congeners to distinct experiences and participates to the emergence of individuality that distinguishes it from others (Bergmüller and Taborsky, 2010; Lathe, 2004). This corresponds to repeatability and consistency of an animal individual behaviour in the face of environmental and social challenges, which translates into different strategies to find food, deal with predators, or compete with conspecifics.

Mice are social vertebrates, living in hierarchical structures of 4 to 12 adult members (Berry and Bronson, 1992; Beery and Kaufer, 2015) that share territorial defence and exhibit a large repertoire of behaviours (e.g. physical exploration, vocal communication, aggression, social recognition, imitation, empathy) that characterize sociability. The social rank of individuals can be determined based on observations of antagonistic interactions, territorial marking, access to limited resources and by precedence behaviours (Zhou *et al*., 2018). The driving forces underlying the emergence of social organization remain largely unknown. Although these include genetic factors, however, the fact that social hierarchy is observed within groups of genetically identical congeners suggests that environmental factors are in play. Among these, the stress response and more specifically glucocorticoids release has been suspected to influence social dominance in a variety of species, although a clear link has yet to be drawn (Sapolsky, 2004; Creel *et al*., 2013).

Hierarchy establishment involves iterative pairwise interactions that have consequences on the behavioural fate of each individual (Cordero and Sandi, 2007;, Timmer and Sandi, 2010). Indeed, specific behavioural patterns emerge in genetically identical mice raised in semi-naturalistic environments (Freund *et al*., 2013;, Hager *et al*., 2014;, Torquet *et al*., 2018), and differences in behavioural traits have been attributed to social ranking in smaller colonies (Wang et *al*., 2011; Larrieu *et al*., 2017). Whether such individual differences pre-exist the formation of the social group is unclear, and the physiological mechanisms implicated in hierarchical segregation remain elusive. Beyond understanding the principles of interindividual behavioural diversity in animals, these questions are also relevant in humans, in a psychopathological context, since social status is recognized as a vulnerability risk factor for psychiatric diseases including mood disorders and addiction (Kessler, 1994; Wilkinson, 1999; Lorant, 2003; Singh-Manoux *et al*., 2005).

The mesocorticolimbic system that encompasses the prefrontal cortex (PFC), the nucleus accumbens (NAcc) and their dopaminergic input from the ventral tegmental area (VTA) could participate to the emergence of social hierarchy and behavioural diversity. This brain system modulates a broad spectrum of behaviours, including motivation and decision-making involved in social context (Gunaydin *et al*., 2014). VTA dopamine neurons activity conditions social avoidance following social defeats (Chaudhury *et al*., 2013; Barik *et al*., 2013). The interaction between stress-evoked release of glucocorticoids and dopamine system is critical for this effect and relies on the activation of glucocorticoid receptors (GR) present in dopamine-sensing areas neurons (Barik *et al*., 2013). Several structures receiving dopaminergic inputs have been recently associated with the emergence of social ranking. Modulating the synaptic efficacy in medial PFC neurons causes individual bidirectional shifts within social ranking (Wang *et al*., 2011), and lower mitochondrial activity within the NAcc is associated to lower social ranking in both rats and mice (Hollis *et al*., 2015; Larrieu *et al*., 2017).

In this study, we examine the segregation of individual behaviours with social status in colonies of four genetically identical male mice (tetrads). We investigated whether pre-existing behavioural and physiological differences shape the social fate of individuals or whether such differences emerge from social life. Finally, we provide evidence for an implication of mesocorticolimbic dopamine system and stress response signalling in the establishment of social hierarchy and individuation of behaviours.

## Results

### Social ranks within tetrads are stable over long periods

We formed colonies of four age and weight matched adult C57BL/6J male mice of six weeks, previously unknown to each other. Two to four weeks later, we analysed the social ranks of animals. We first used a precedence test based on encounters within a plastic tube between each possible congener pairs among a tetrad. It allows to identify lower ranked individuals as they come out of the tube walking backward (Wang *et al*., 2011) (Fig. 1a). The six possible pairwise combinations of individuals from a tetrad were tested three consecutive times a day, and the one with the highest number of forward exits was classified as higher ranked. We tested each tetrad daily, for at least six days, until the highest (rank 1 rated R1) and the lowest ranks (rank 4 rated R4) were stable over 3 consecutive days.

**Figure 1.**
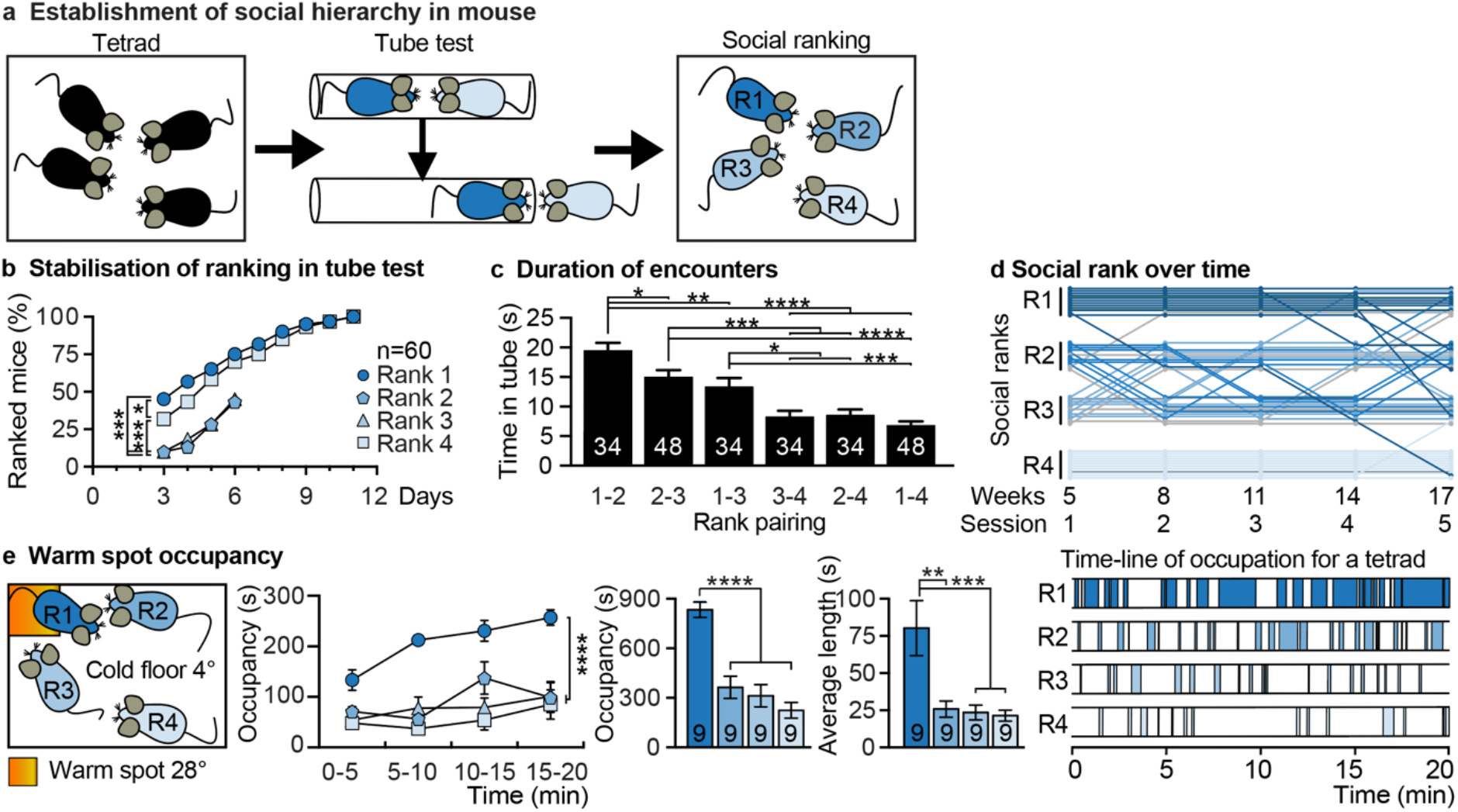
Social hierarchy establishment and stability in mice. **a**, Design of social hierarchy establishment and analysis. Unfamiliar male mice were grouped by four. After 3 to 4 weeks, their social rank was determined by a precedence test (tube-test), and further tested four times until week 17. **b**, Rapidity of rank identification in the tube-test. The cumulated percentage of stable ranked individuals for each rank is pictured for each day of tube-test (n=60 tetrads). The data for ranks 2 and 3 are indicated for days 3 to 6 since rank identification was stopped when ranks 1 and 4 were stable. Gehan-Breslow-Wilcoxon Test. **c**, Mean duration of the confrontation in the tube-test performed during the three last days when ranks were stable Each possible rank combination is pictured. The numbers of tetrads that have been analysed are indicated for each rank combination; n=48 or 34 in cases of unstable ranks 2 and 3. Wilcoxon rank sum test. **d**, Social ranks were stable from week 5 and for over three months for most animals. The dynamic of social ranking in the tube-test is pictured for a set of 12 tetrads. Each line depicts an individual mouse, its position within its social rank pool indicates the tetrad which it belongs to. Different blue intensities indicate the rank defined at the first tube-test session. The 6 individuals of ranks 2 and 3 that did not reach stability at the end of the first session are pictured with by grey lines. **e**, Left : representation of the warm spot test. The position of the warm spot is pictured by an orange box. Middle : time course occupancy of the warm spot, total occupancy, and average length occupancy by differently ranked individuals (n=9 tetrads). Right : representative occupancy periods of the warm spot by individuals of a tetrad. *p < 0.05; **p < 0.01; ***p < 0.001. Error bars, ± SEM.

Among 60 tetrad colonies, the stability criterion was reached faster for the extreme ranks (Fig. 1b). Half of R1 and R4 mice were already having a stable status on days 3 and 5, respectively. All of them were stable after 12 days whereas a quarter of ranks 2 (R2) and rank 3 (R3) animals were not. As observed by Wang *et al*., 2011, the rank of individuals conditioned the duration of contests. Confrontations between R1 and R2 individuals lasted for an average of 18 seconds whereas confrontations involving R4 lasted twice less (Fig. 1c).

Once established, social ranking was stable over long periods. Fig. 1d pictures social fate of individuals from 12 tetrads repeatedly assessed through 5 sessions during a period of 3 months. This is particularly true for R4 individuals (Fig. 1d, lighter blue lines) as 17 weeks later, eleven out of twelve mice remained at the lowest rank. Among initially highest ranked individuals, ten and seven, out of twelve, kept the same ranking, 14 and 17 weeks later, respectively (Fig. 1d, darker blue lines). One progressively decreased ranking, to end in R4, one ended in R3 and three in R2. Animals with initial intermediate ranks displayed the highest switching in rankings but 19 out of 24 still ended by reaching an intermediate rank (Fig. 1d, medium blue and grey lines). During this period of time, mice were regularly weighted, and no correlation between social rank and weight evolution was found (data not shown).

To validate precedence behaviour as a reliable proxy for social ranking we quantified other expressions of social dominance, such as access to shared resources or territoriality. Higher ranked individuals in the tube-test showed significantly longer total occupancy of a small warm spot within a cold cage during the 20 minutes of the test (Fig. 1e) compared to their three other cage-mates, with 3 to 4 times longer episodes. Urine marking patterns were collected on absorbent paper from a box occupied by R1 and R4 individuals separated by a transparent and perforated wall for 2 hours and visualized under U.V. light (extended data Fig. 1a). 17 out of 23 top-ranked individuals in the tube-test also showed dominant urine marking patterns, in either the number of marks or their cumulated area, when compared with lower ranked congeners.

### Social rank correlates with behavioural differences

We compared behaviour between higher (R1) and lower (R4) socially ranked individuals. We did not see differences neither in locomotor activity, measured in an open-field (data not shown), nor in stress-coping, measured by quantifying immobility and escape behaviours in the forced-swim test for two consecutive days (Fig. 2a).

**Figure 2.**
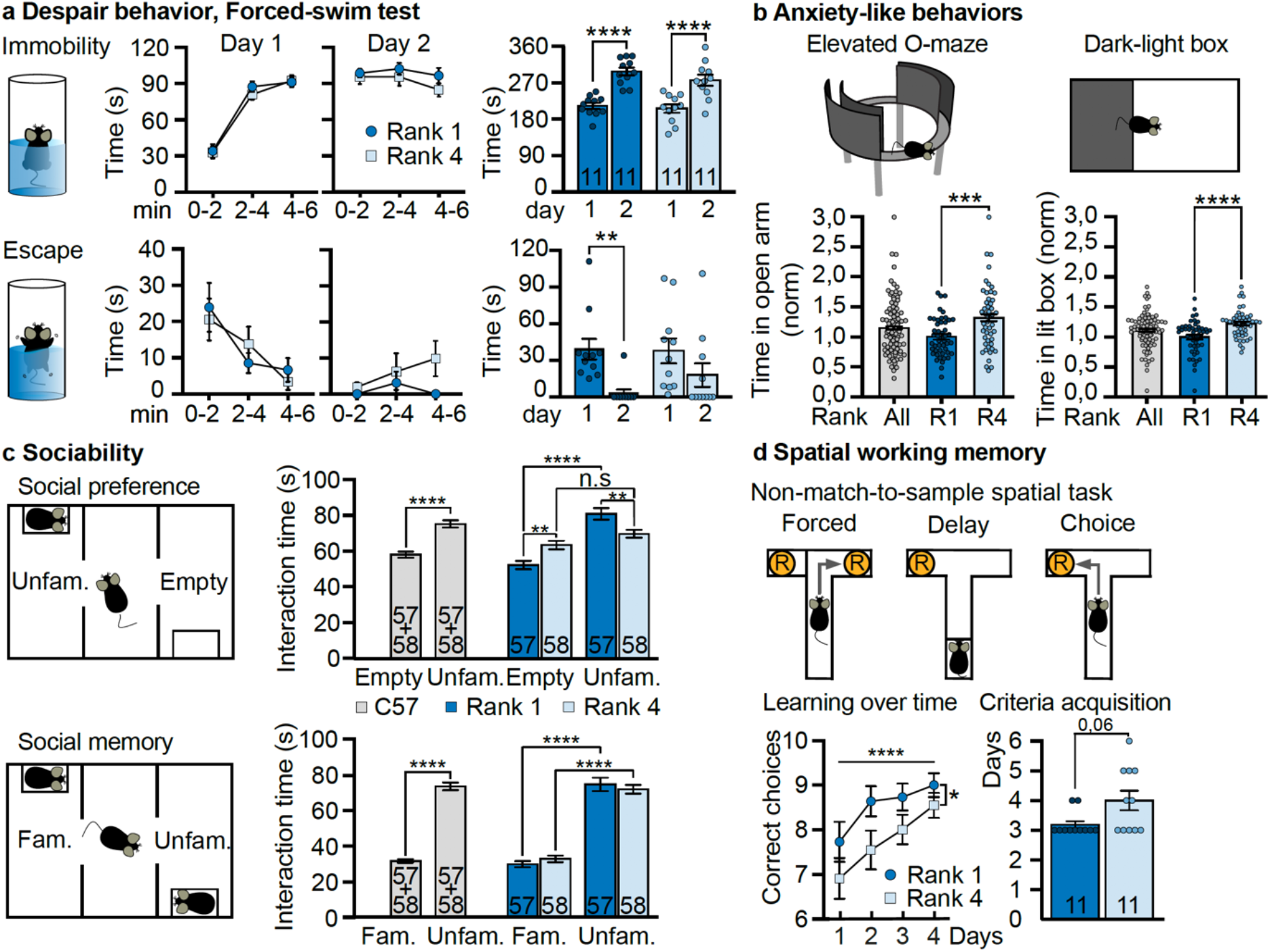
Differences in social rank correlates with differences in behaviour in genetically identical mice. **a**, Highest and lowest ranked animals display similar depression-like behaviour in the forced-swim test. The length of immobility in tepid water is presented by 2 minutes periods (upper panel, left) and for the 6 minutes of the test (upper panel, right) for R1 and R4 mice (n=11 per group). Results obtained on the first day (Day 1) and 24 h later (Day 2) are pictured. The lower panels present, in the same way, the quantification of escape behaviour of the same individuals. Error bars, ± SEM. **b**, Rank 1 individuals display increased anxiety-like behaviours. The experimental setups are pictured. The time spent for all C57B/L6 (n=96), Rank 1 (n=48, dark blue) and Rank 4 (n=48, light blue) individuals in the open-section of an elevated O-maze and in the lit compartment of a dark-light box are pictured. Time values were normalized from the R1 means. Respectively, t94=3.55, ***p<0.001 and t94=4.213, ****p<0.001, unpaired t-tests, two-tailed. Error bars, ± SEM. **c**, Highest ranked mice display increased sociability but a social memory similar to that of lowest ranked individuals. The three-chambers test is depicted (left). The duration of time spent interacting with an empty box (Empty) and with a box containing an unfamiliar mouse (Unfam.) are pictured in the upper row for C57B/L6 (grey, n=115), R1 (dark blue, n=57) and R4 (light blue, n=58) individuals. The duration of time spent interacting with a box containing a familiar mouse (Fam.) *vs* an unfamiliar one (Unfam.) are represented in the lower row. Upper row: t114=6.012, **** p<0.0001, unpaired t-test, two-tailed. Right graph: effect of interaction, **** p<0.0001, F_(1,113)_=17.07; effect of social cue, ****p<0.0001, F_(1,113)_=41.7; no effect of social rank, p=0.97, F_(1,113)_=0.002. Empty box: R1 *vs* empty box: R4, **p<0.01; Social cue R1 *vs* social cue R4, **p<0.01; Empty box *vs* social cue for R1 mice, **** p<0.0001. Empty box *vs* social cue for R4 mice, p=0.20. Two-way mixed ANOVA, Bonferroni’s test. Error bars, ± SEM. Lower row : t114=15.19, **** p<0.0001, unpaired t-test, two-tailed. Right graph: no effect of interaction, p=0.30, F_(1,113)_=1.097; effect of familiarity, **** p<0.0001, F_(1,113)_=231.1; no effect of social rank, p=0.003, F_(1,113)_=0.97. R1: familiar *vs* unfamiliar, ****p<0.0001; R4: familiar *vs* unfamiliar, ****p<0.0001. Two-way mixed ANOVA, Bonferroni’s test. Error bars, ± SEM. **d**, Rank 1 individuals display better performances in the Non-match-to-sample-spatial task, a spatial working memory task. Upper row illustrates the task design. The learning curve of 11 mice from both ranks, indicates the progression of correct choices over the days (lower row, left). The number of days required to reach the learning criterion is indicated (mean and individual scores, right). Effect of time, ****p<0.0001, F_(3,60)_=7.87; effect of social rank, *P<0.05, F_(1,20)_=4.85; no effect of interaction, p=0.78, F_(3,60)_=0.36. Two-way mixed ANOVA. Right panel indicates for each rank the average number of days required to acquire the criterion. U=34.5, p=0.056 Mann-Whitney U test, two tailed. Error bars, ± SEM.

In contrast, anxiety-like behaviour and sociability markedly differed between social ranks, highest ranked individuals being more anxious and more sociable than lower ranked ones. We quantified anxiety-like behaviour approach-avoidance conflict tests based on the mouse innate avoidance of open and lit spaces (Fig. 2b). When measuring anxiety of isogenic mice, high interindividual differences are classically observed, as in Fig. 2b (grey dots). Considering individual social ranks allow to stratify the population into two distinct groups with regards to anxiety-like behaviours. R1 individuals, more anxious, spent significantly less time in the open section of an elevated O-maze (Fig. 2b, left dark blue), as well as in the lit compartment of a dark-light box (Fig. 2b, right dark blue). We also compared sociability and social memory between R1 and R4 mice in a three-chamber test. As expected, C57BL/6 mice display a marked preference for a social stimulus (Fig. 2c, social preference, grey bars). Stratification of the results taking into account the social rank of individuals shows that only the R1 individuals, and not the R4, displayed social preference (Fig. 2c, social preference, right panel, dark and light blue bars, respectively). However, social rank does not affect social memory or social novelty. Mice of both ranks have a similar natural preference for interacting with an unfamiliar conspecific *vs* a familiar one (Fig. 2c, lower panel).

During manipulations of the tetrads, R4 mice frequently showed aggressive behaviours towards their cage-mates. We performed a resident-intruder test, repeated for two consecutive days with individuals from 10 tetrads. From half of them, at least one R1 or R4 mice attacked the intruder. For these pairs, we quantified the interactions with the intruder (extended data Fig 2). On the first day, R4 individuals displayed significantly more aggressive behaviours, including clinch attacks, lateral threats, chases, upright postures and bite attempts whereas R1 individuals had more prosocial behaviour (grooming of the intruder). On the second day this difference was more pronounced. Whereas none of the R1 individuals attacked the intruder, all R4 mice did within the first 130 seconds and did significantly more rattling.

We then addressed whether social ranking could also affect cognitive abilities. We studied spatial working memory in a non-match-to-sample T-maze task (Fig. 2d, upper panel). In this task, mice are placed within a T-Maze and can access a reward placed into the unique open arm (forced phase). They are required to retain a memory trace of a recently sampled maze location during a delay period (delay phase) and then prompted to select the opposite location in order to receive a reward (choice phase). Each mouse was tested 10 times a day, and the learning criterion was defined as a minimum of 7 correct choices for 3 consecutive days. Although both groups of mice learned the task, R1 individuals did it significantly faster compared to R4 (Fig. 2d, lower panel).

### Differences in sociability but not anxiety-like behaviours pre-exist to social rank establishment

The behavioural differences between ranks could emerge from initially similar mice as a consequence of social adaptation within members of a tetrad. Alternatively, these differences could pre-exist before their gathering, and shape individual social ranking trajectories. To address this question, we compared behaviours of individuals before grouping them in tetrads, and after the formation of the social colony. As mentioned before, anxiety-like behaviours were markedly enhanced for highest ranked individuals compared to lowest ones. This behavioural difference seems to emerge from social organization since no difference was observed between future R1 and R4 mice, before they were pooled together, in elevated O-maze and in dark-light tests. The time spent in open arms and in the lit compartment were similar for these two groups, as well as for the future ranks 2 and 3 (Fig. 3b). In contrast differences in sociability seem to pre-exist to life in colony. Future R1 mice have already a marked interest for social interactions before social life in tetrads, similar to that observed once the tetrad is formed (Fig. 3c, dark blue bars, left panel), whereas future R4 have not. Interestingly, intermediate ranks have an intermediate phenotype with a significant but lower preference. R1 and R4 mice did not present differences in despair-like behaviour, thus as expected we did not observe any differences between future R1 and R4 (Fig. 3a) individuals.

**Figure 3.**
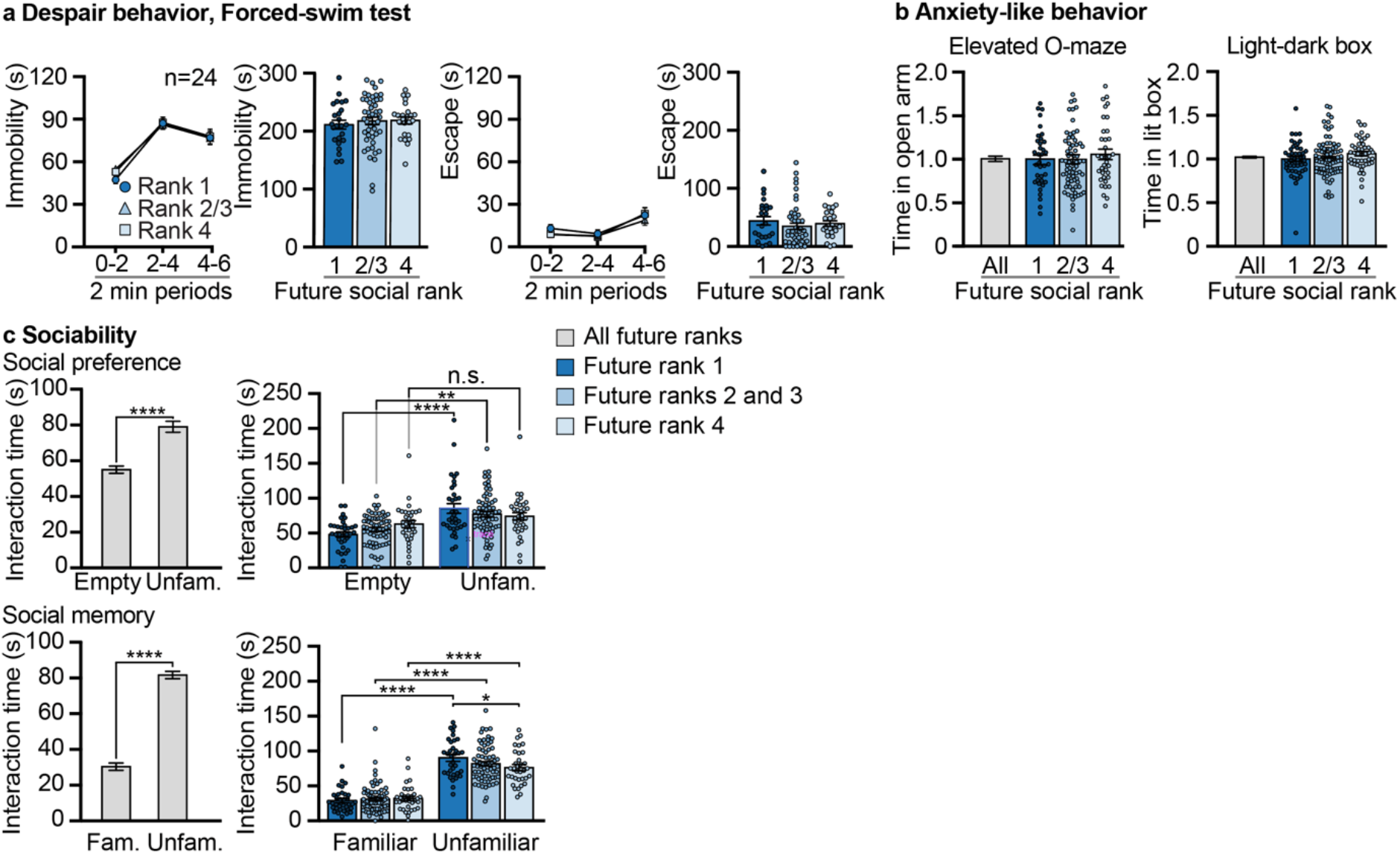
Difference in anxiety-like behaviour emerge for social life but difference in sociability is a trait pre-existing to social life. **a**, Future rank 1 and rank 4 individuals have similar despair behaviour in a forced swim test. The time length of immobility and escape are pictured for each periods of two minutes (lines), and for the six minutes of the test (bars). For future R1 (n=24), R4 (n=24), and R2/3 (n=48) (dark, medium and light blue, respectively) Error bars, ± SEM. **b**, Future rank 1 and rank 4 individuals display similar anxiety-like behaviours. The time spent to explore the open segments of an elevated O-maze and the lit compartment of a dark-light box are pictured for all C57BL/6 mice (grey bars, n=144 and n=192, respectively), and among them the future R1 (dark blue bars, n=36 and n=48, respectively), the future ranks 2 and 3 (medium blue bars, n=72 and n=95, respectively), and the future R4 (light blue bars, n=36 and n=48, respectively). For the O-maze, one dot out of scale (3.25) was omitted for ranks 2/3. Scores are normalized from the R1 means. Error bars, ± SEM. **c**, Social behaviour. Social preference (left), Future rank 4 mice did not display social preference unlike future R1 individuals. Duration of interactions with an empty box *vs* a box containing an unfamiliar (Unfam.) congener is pictured for 136 mice C57BL/6 mice from tetrads (grey bars) and, among them, the future R1 (dark blue bars, n=34), R2/3 (medium blue bars, n=68) and R4 (light blue bars, n=34). Left: t135=5.435, ****p<0.0001, unpaired t-test, two tailed. Right: no effect of interaction, p=0.09, F_(2,133)_=2.41; effect of social cue, ****p<0.0001, F_(1,133)_ 27.55; no effect of social rank, p=0.77, F_(2,133)_=0.28. Empty box *vs* social cue for R1 mice, ****p<0.0001; empty box *vs* social cue for R2/3 mice, **p≤0.001. Two-way mixed ANOVA, Bonferroni’s test. Error bars, ± SEM. Social memory. The time length interaction with the box containing a familiar (Fam.) social cue *vs* a box containing an unfamiliar (Unfam.) mouse is pictured. Left: t135=17.71, ****p<0.0001, unpaired t-test, two tailed. Right: no effect of interaction, p=0.01, F_(2,133)_=2.31; effect of familiarity, ****p<0.0001, F_(1,133)_=291.6; no effect of social rank, p=0.34, F_(2,133)_=1.10. Familiar *vs* unfamiliar for R1 mice, ****p<0.0001; Familiar *vs* unfamiliar for R2/3 mice, ****p<0.0001; Familiar *vs* unfamiliar for R4 mice, ****p<0.0001. Unfamiliar social cue: R1 *vs* R4, *p<0.05. Two-way mixed ANOVA, Bonferroni’s test. Error bars, ± SEM.

### Social rank conditions sensitivity to some preclinical models of psychopathologies

In human, low social status, is associated with reduced life span (Stringhini *et al*., 2017) and negative health consequences including chronic cardiovascular, immune, metabolic and psychiatric disorders). In a variety of social vertebrate species, associations between social rank and health outcomes have been documented (Sapolsky, 2005). We investigated whether highest and lowest ranked individuals within tetrads would respond distinctly to two established preclinical models of mental disorders in mice.

The locomotor sensitization to cocaine is a gradual and enduring facilitation of locomotor activity promoted by repeated cocaine exposure, believed to reflect the reinforcing effects of abused drugs (Robinson and Berridge, 2000). Five days of daily cocaine injections (10 mg.kg^−1^) led to a gradual increase of locomotion during the hour following the drug administration in both R1 and R4 mice (Fig. 4a). This increased locomotor response was maintained after a withdrawal period of 6 days as assessed after a challenge injection (Fig. 4a, day 12). While this result clearly shows that both R1 and R4 individuals exhibit significant locomotor sensitization to cocaine, R4 individuals appeared more sensitive. Indeed, R4 mice showed enhanced responses during the five consecutive cocaine injection days, and an even more pronounced hypersensitivity during the challenge day (Fig 4a, right panel).

**Figure 4.**
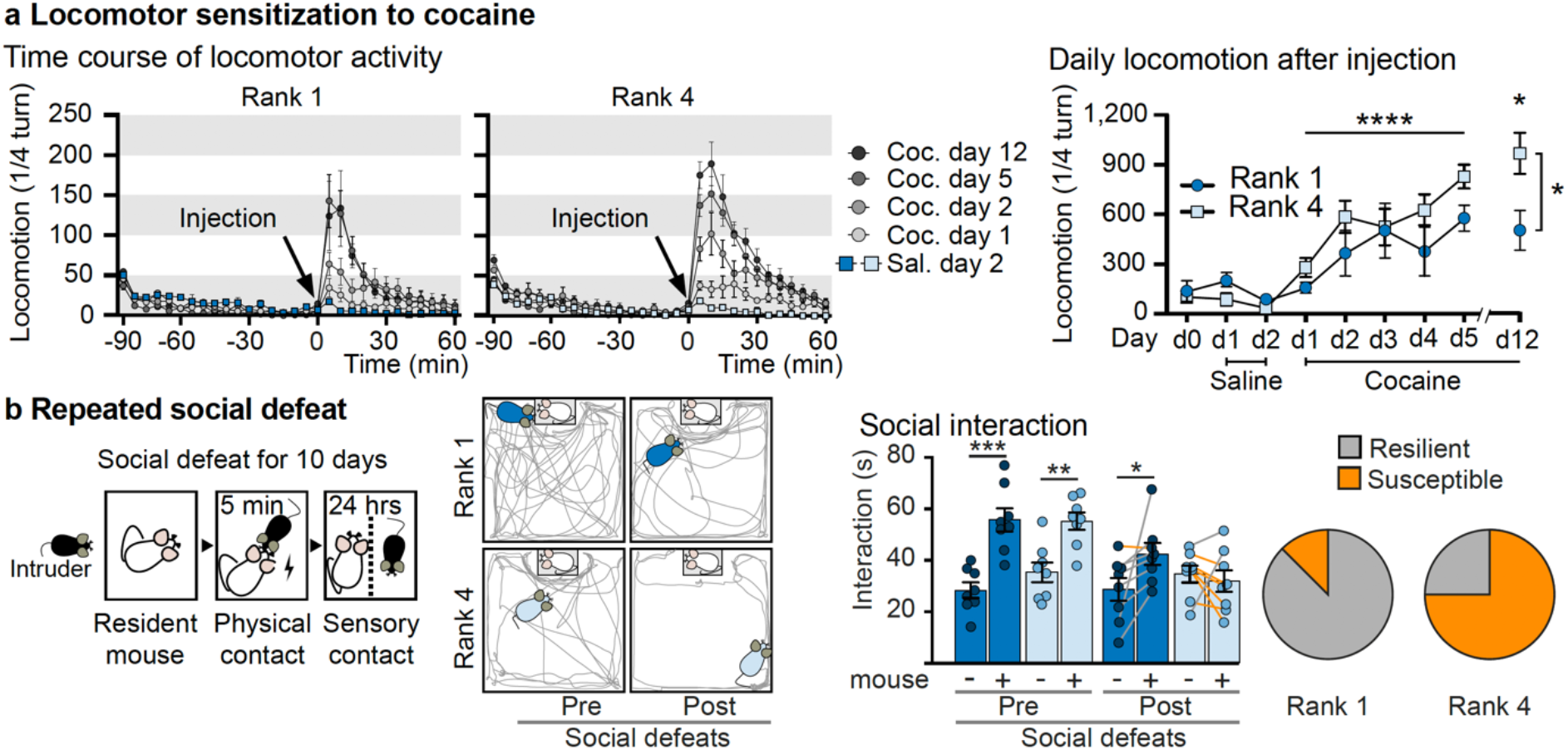
Social rank correlates with differences in preclinical models of behavioural disorders. **a**, Locomotor sensitization to cocaine. Left, time course of Rank 1 (left panel, n=8) and Rank 4 (middle panel, n=9) individuals locomotion expressed as ¼ turn within a circular corridor for indicated sessions. Time 0 correspond to the injection of cocaine (Coc, 10 mg kg^−1^) or saline (Sal). Right panel, cumulated locomotor activity of R1 and R4 individuals (dark and light blue, respectively) for 1 hour following habituation (day 0), saline (days 1 and 2) and cocaine (days 1 to 5 and a challenge on day 12) injections. Effect of time, **** p<0.0001, F_(5,75)_=7.55; effect of social rank, *p<0.05, F_(1,15)_=5.376; no effect of interaction, p=0.32, F_(5,75)_=1.198. Coc d12 R1 *vs* Coc d12 R4, *p<0.05. Two-way mixed ANOVA, Bonferroni’s test. Error bars, ± SEM. **b**, Depression-like behaviour induced by repeated social defeats. Left, protocol design. Middle left, representation of the open field in which social interactions were measured. The position of the box containing a CD1 mouse is indicated. Representative trajectories of R1 and R4 individuals before and after repeated social defeats are pictured. Middle right panel, R1 (dark blue, n=8) and R4 (light blue, n=8) interaction time with an empty box (mouse -) or a CD1 mouse (mouse +), before and after repeated social defeat. Right panel, susceptible individuals, developing social aversion are indicated with orange lines, resilient ones with grey ones. Right, representation of susceptible (orange) and resilient (grey) individuals among R1 and R4 males. Pre-social defeats: effect of social cue, **** p<0.0001, F_(1,14)_=41.2; no effect of social rank, p=0.39, F_(1,14)_=0.76; no effect of interaction, p=0.33, F_(1,14)_=1.04. Empty box *vs* social cue for R1 mice, *** p<0.001; empty box *vs* social cue for R4 mice, **p<0.01. Post-social defeats: no effect of social cue, p=0.19, F_(1,14)_=1.89; no effect of social rank, p=0.60, F_(1,14)_=0.29; effect of interaction, *p<0.05, F_(1,14)_=4.95. Empty box *vs* social cue for R1 mice, *p<0.05. Two-way mixed ANOVA, Bonferroni’s test. Error bars, ± SEM.

Repeated social defeats is a well-validated mouse model of depression, marked by a lasting social aversion, which allows to distinguish animals that exhibit depressive-like symptoms (susceptible) from those which are resilient to stress (Krishnan *et al*., 2007). Highest and lowest ranked mice from eight tetrads were daily subjected to 5 minutes social defeats for 10 consecutive days by an unfamiliar aggressor and remain in sensory (but physical) contacts for the rest of the day (Fig. 4b, left panel). We quantified in an open field the time of interaction with an empty plastic box *vs* a box containing an unfamiliar male mouse, before and after social defeat (Fig 4b middle panel). After social defeats, 7 out of the 16 mice challenged developed a social aversion (Fig. 4b right panel, orange lines). Among them only one was a R1 individual. In other words, majority of R1 individuals (87.5%) were resilient whereas only 25% of R4 ones were.

### Dopamine neurons activity in the ventral tegmental area and glucocorticoid receptor in dopamine-sensing neurons modulate social rank attainment

Stress response and glucocorticoids release is a major risk factor for psychopathologies. We and other have shown that the interplay between stress response and the dopamine reward pathway plays a major role in stress-induced depressive behaviour and responses to addictive drugs (Ambroggi *et al*., 2009; Barik *et al*., 2010; Barik *et al*., 2013). Several lines of evidence suggest that both systems could play a significant role in rank establishment. We first compared dopamine utilization in R1 and R4 individuals in key brain regions innervated by midbrain dopamine neurons.

We measured in the caudate putamen (CPu), the NAcc, and the PFC the tissue contents of dopamine (DA) and 3,4-dihydroxyphenylacetic acid (DOPAC), a metabolite produced following dopamine reuptake. The DOPAC/DA ratio thus gives an index of dopamine utilization (Fig. 5a left graph). We did not observe any difference in the CPu, a structure mostly innervated by dopamine cells located in the substantia nigra pars compacta (SNc). In the NAcc, we noted a trend towards a decreased dopamine utilization in R1 individuals when compared to R4 ones, although the difference did not reach significance (Fig. 5a). In striking contrasts, in the PFC, R1 individuals displayed a marked increase of dopamine release. Overall, we showed a dopamine utilization characterized by a stronger PFC/NAcc ratio in R1 mice compared to R4 ones (Fig. 5a right graph). Since both the NAcc and the PFC are mainly innervated by dopamine cells located in the VTA, we investigated whether differences could exist in their activity between R1 and R4 individuals. We performed juxtacellular single-unit recordings in anesthetized mice. The analysis of 186 neurons from 10 R1 mice and 157 neurons from 10 R4 mice revealed that whereas the frequency of spontaneous firing was similar in both ranks (Fig. 5b, left graph), the percentage of spikes within bursts was significantly lower in R1 individuals (Fig. 5b, right graph).

**Figure 5.**
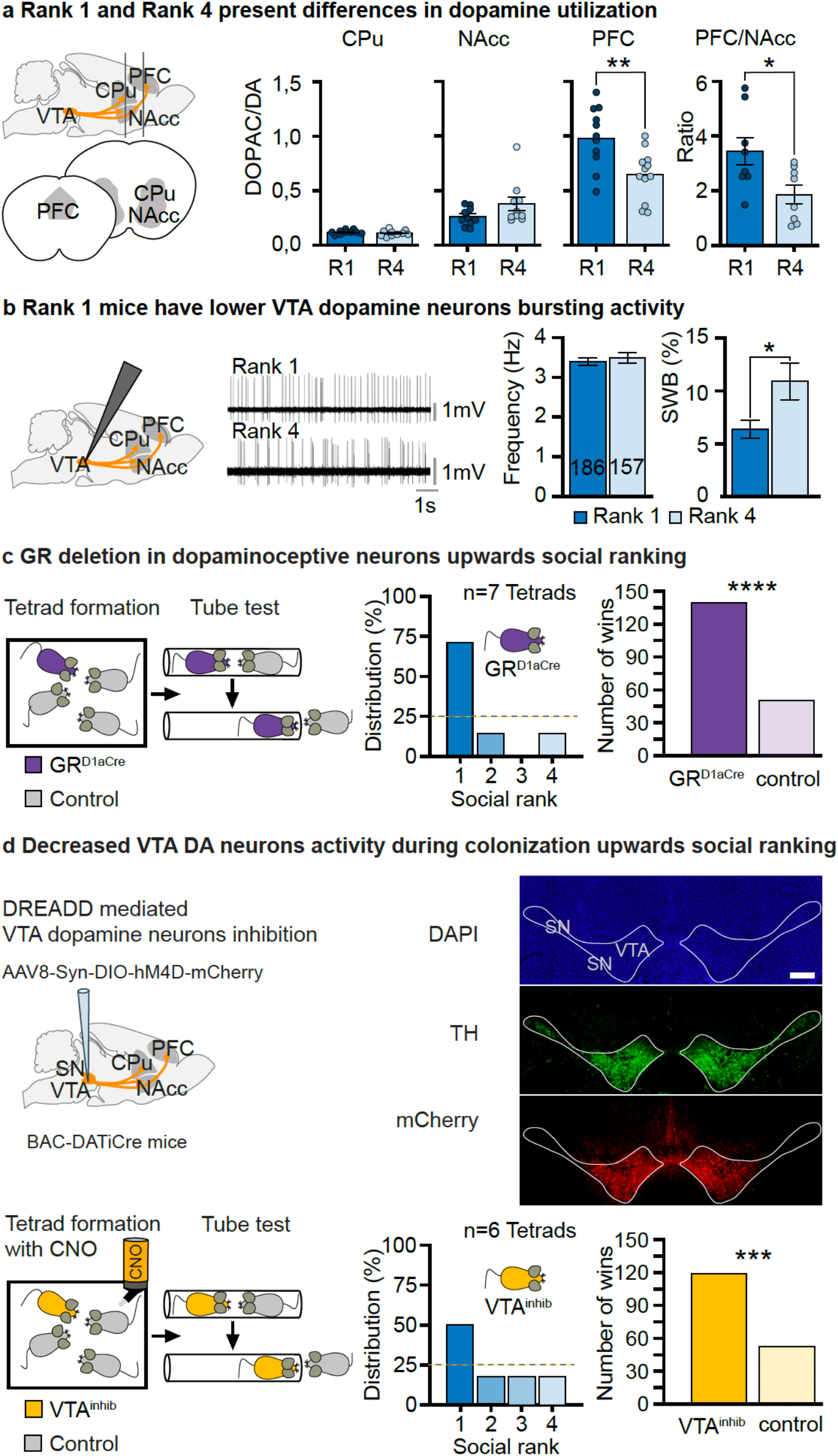
Dopamine neurons activity in the ventral tegmental area and Glucocorticoid Receptor in dopaminoceptive neurons modulate social rank attainment. **a**, Differences in dopamine utilization between Rank 1 and Rank 4 males. Left, sagittal representation sketches the section lines for tissue punches along the mesocorticolimbic pathway (bottom, coronal view). Dopamine utilization was estimated measuring the ratio DOPAC/DA in the caudate putamen (CPu, n=10, 5 mice), the nucleus accumbens (NAcc, n=10, 5 mice) and the prefrontal cortex (PFC, n=12, 6 mice) of both R1 and R4 mice. The PFC/NAcc ratio was calculated for 4 mice (n=8). For PFC, t22=3.256, **p<0.01, For PFC/NAcc t14=2.51, *p<0.05, unpaired t-tests, two-tailed. n represents the number of hemispheres for each group. Error bars, ± SEM. VTA: ventral tegmental area. **b**, Rank 1 individuals have lower VTA dopamine neurons bursting activity. Schematic representation of electrode positioning (left). Representative traces of recording for individuals of each rank (middle). Mean frequency (Hz) and percentage of spikes within bursts (SWB) of dopamine cells basal firing mice belonging to R1 (n=186, 10 individuals) and R4 (n=157, 10 individuals) (right). For SWB data: t341=2.362, p*=0.02, unpaired t-test, two-tailed. Error bars, ±SEM. **c**, GR deletion in dopaminoceptive neurons promotes higher social ranking in tetrads. Left, tetrads were constituted with one mutant (GR^D1aCre^) and three control (GR^lox/lox^) mice, for three weeks. The middle graph indicates the percentage of GR^D1aCre^ mice reaching each rank, among the 7 tetrads tested. The right graph pictures the number of won tube test contests between GR^D1aCre^ and control mice, out of 189 encounters during the 3 last days of tube test sessions. Fischer’s exact test, two-tailed, ****p<0.0001, Error bars, ± SEM. **d**, Decreased VTA dopaminergic neurons activity during colonization upwards social ranking. Upper row, schematic representation of the DREADD expressing AAV8 particles injection site (left). Representative analysis of injection site (right). Upper panel, DAPI staining, VTA: ventral tegmental area, SN: substantia nigra. Bar: 250 μm. Middle panel, immunofluorescence staining detecting Tyrosine Hydroxylase (TH) expression. Lower panel, immunofluorescence staining detecting mCherry expression. Lower row, tetrads were constituted with one mouse expressing CNO-dependent hM4D (VTA^inhib^) and three expressing GFP in the VTA (control) mice. CNO was permanently delivered in drinking water. After two weeks, social hierarchy was determined. The middle graph indicates the percentage of VTA^inhib^ mice reaching each possible rank, among the 6 tetrads tested. The right graph pictures the total number of won tube test contests between VTA^inhib^ and control mice, out of 171 encounters during the 3 last days of tube test sessions. Fischer’s exact test, two-tailed, **** p<0.0001, *** p<0.001 Error bars, ± SEM.

### Reduced activity of VTA dopamine neurons facilitates higher ranking

Glucocorticoid receptor (GR) gene inactivation in dopamine innervated areas results in lower VTA dopamine neurons activity, similar to that observed in R1 mice (Ambroggi *et al*., 2009; Barik *et al*., 2013). Further, we showed that it facilitates resilience to social defeat by preventing the appearance of social withdrawal and reduces behavioural responses to cocaine (Ambroggi *et al*., 2009; Barik *et al*., 2010; Barik *et al*., 2013). We thus examined whether GR signalling within the dopamine reward pathway could influence social ranking in tetrads. We grouped one adult GR^D1aCre^ mice with three unfamiliar control (GR^loxP/loxP^) individuals (Fig 5c, left graph) and assessed their social rank in tube-test two weeks later. Within 5 tetrads out of 7, GR^D1aCre^ mice ended on the highest rank, in one it was ranked second and in one fourth (Fig 5c, middle panel). Analysing tube-test individual contests during the last three days of ranking determination, we observed that mutant mice have a significantly higher probability to win. Out of 189 contests involving a mutant and a control mouse, mutant ones won 139.

These results raise the possibility that stress-response might influence social fate by impacting on the mesolimbic dopamine neurons. To test the causality between low VTA dopamine neurons activity and higher social rank, we assessed the impact of inhibiting this cell population on social hierarchy. To do so, we expressed a modified human muscarinic receptor Gi coupled hM4D specifically in VTA dopamine neurons, using an adeno-associated virus expression system (AAV8-hSyn-DIO-hM4D(Gi)-mCherry), stereotactically injected into the VTA of mice expressing the Cre recombinase in dopamine neurons (BAC-DATiCre-Turiault *et al*., 2007-, VTA^inhib^ mice, Fig. 5d). The hM4D receptor is activated solely by a pharmacologically inert compound, clozapine-N-oxide (CNO). Upon CNO activation, hM4D hyperpolarizes neurons through a Gi protein mediated activation of inward-rectifying potassium channels (Armbruster *et al*., 2007). Control animals were C57B/L6 mice similarly injected with AAV8-hSyn-GFP particles. Tetrads composed of one VTA^inhib^ mouse and three control mice were formed and permanently treated with CNO in drinking water (10 mg L^−1^, Fig. 5d left). After analysis of the social hierarchy in these tetrads, we observed that permanently reducing the activity of VTA dopamine neurons increased the probability of the individual to reach the highest rank, (Fig. 5d middle) and significantly increased the number of tube test wins. Out of 171 contests involving a VTA^inhib^ and a control mouse, VTA^inhib^ ones won 118 (Fig. 5d right). In another experimental design, we tested the effect of acute dopamine neurons inhibition on tube test performance. We found that a single injection of CNO (1 mg kg^−1^) 30 minutes before tube test contests changed neither the rank of VTA^inhib^ mice, nor their probability to win a contest in already established tetrads (extended data Fig. 5, Day 1). However, after several days of CNO injections, their probability to win a contest seemed to increase (from 49,2 ± 15,9 % on day 1 to 67,5 ± 14,7 % on day 5, n=7 individuals), strengthening the idea that VTA dopamine neurons activity is rather important for the process of hierarchy establishment than for punctual tube test performance (extended data Fig. 5).

## Discussion

Within a few days, genetically identical mice living in small groups of four individuals establish a social organization. The social hierarchy can be determined observing differential precedence in displacements as in tube-test (Wang *et al*., 2011; Larrieu *et al*., 2017) and differential access to resources, as a warm spot in a cold environment. The attained social rank is stable over month periods, with limited switch between ranks within a tetrad. These switches are almost absent for the lowest ranked individuals and rarely observed for the highest ones. They are frequent for the two intermediate ranks that also take a longer time to stabilize during the first tube-test session. Behavioural analyses usually present an important interindividual variability and social ranking might at least partly account for this phenomenon. We showed, in agreement with Larrieu *et al*. (2017) that used the same ranking methods, that highest ranked mice exhibit indeed higher anxiety-like behaviours and increased social interactions. Varholick *et al*. (2018) did not however observe this correlation. This discrepancy may rely on the limited number of animals tested, or on the approach they chose to identify ranking, with sparser tube tests, performed once a week for three weeks. Few other studies that used other criteria to identify dominant individuals, such as aggressiveness, have been carried out leading-to conflicting results. As such, Hilakivi *et al*. (1989) reported no difference whereas Ferrari *et al*. (1998) saw an increased anxiety for dominant individuals. Similarly, the high dispersion of individual interaction time with a congener during sociability tests in isogenic mice can also be in part explained by their social rank. Highest ranked ones are indeed more sociable, in agreement with Kunkel and Wang (2018). This association of high anxiety and high sociability is surprising as in both humans and rodents, low anxiety is usually paired with increased sociability (Allsop et al., 2014; Beery and Kaufer, 2015). For instance, oxytocin facilitates social behaviors and has well-known anxiolytic properties (Insel, 2010). In the same line, optogenetic stimulation of basolateral amygdala to ventral hippocampus circuit facilitates anxiety and impairs social interaction (Felix-Ortiz *et al*., 2013; Felix-Ortiz and Tye, 2014). This associative rule is nevertheless not systematic. Similar to our observation in top ranked individuals, vasopressin promotes social behaviour and is also anxiogenic (Bielsky *et al*., 2004).

A central but poorly explored question is whether the emergence of social ranks precedes the appearance of specific individual behavioural traits, or whether pre-existing individual differences channel the social status trajectory of an individual. Our study indicates that both situations occur. The anxiety of highest ranked animals clearly emerged following social life since no differences existed prior to rank establishment. A similar observation was made in outbred Swiss mice, housed in dyads and ranked upon their aggressiveness (Hilakivi-Clarke and Lister, 1992). On the contrary, in rats, a study showed that a high level of anxiety is a predisposing factor for social submission (Hollis *et al*., 2015). In our study, while the increased anxiety seems to be the consequence of social ranking, the difference in sociability clearly preexists to the formation of the ranks within the colonies. The origin of this individual difference in behaviour most likely arises due to previous breeding and social housing conditions. It could have emerged in the first colony in which these animals were grouped. It could also occur from differences that appeared early, before weaning, since a study suggested that maternal care could shape adult social behaviour (Starr-Phillips and Beery, 2014).

Several studies point at the mesocorticolimbic system as a potential substrate for social ranking. In the NAcc, low mitochondrial activity has been causally linked with lower rank in dyadic contests (Hollis *et al*., 2015). Also, increased activity and higher strength of excitatory inputs to the PFC layer V has been linked with higher social ranking (Wang *et al*., 2011). Our study shows that VTA dopamine neurons exhibit differential activity depending on the rank, with a marked reduction of the bursting activity in R1 individuals. Several studies suggest a role for dopamine in social ranking from insects to mammals (Yamaguchi *et al*., 2015). In ants, brain dopamine concentration is higher in socially dominant individuals (Penick *et al*., 2014; Okada *et al*., 2015). In birds and lizards, increased levels of dopamine in striatal structure have been observed in higher ranked individuals (McIntyre and Chew, 1983; Korzan *et al*., 2006). In line with our results, reduced levels of dopamine have been shown in the NAcc of dominant rats (Jupp *et al*., 2016).

Genetic evidence also sustains a link between dopaminergic neurotransmission and social status. Dopamine transporter gene is essential for sensing dopamine release and dopaminergic neurotransmission. Its inactivation in mice disorganizes social colonies (Rodriguiz *et al*., 2004), and genetic variants of the DAT gene are associated with social dominance in macaques (Miller-Butterworth *et al*., 2007). Imaging studies in humans and nonhuman primates showed an enhanced availability of the striatal D2 receptor for individuals with dominant status. This could result from either higher level of D2 receptors or lower dopamine release (Nader *et al*., 2012; Cervenka *et al*., 2010; Martinez *et al*., 2010). Neuropharmacological approaches also suggest a role for dopamine signalling in social ranking but the differences in strategies deployed (e.g. systemic *vs* local striatal injections in the NAcc) do not allow clear interpretation. Systemic administration of D2 receptor antagonist reduced social dominance in both mice and monkeys (Yamaguchi *et al*., 2017a) whereas local injection into the NAcc of an agonist did not have an effect in rats(van der Kooij *et al*., 2018). Similar experiments with a D1 receptor antagonist facilitated or did not modify social dominance in mice and monkeys(Yamaguchi *et al*., 2017b) whereas local injection into the NAcc of an agonist increased dominance in rats (van der Kooij *et al*., 2018). Interestingly, changes in VTA dopamine cells activity are observed during the emergence of behavioural categories occurring within groups of dozens of mice living in complex semi-naturalistic environments (Torquet *et al*., 2018). It would be interesting to study social ranking between these categories using precedence tests such as the tube test. The decreased firing in R1 mice was associated with a trend of decreased dopamine release in the NAcc as measured by the ratio DOPAC/DOPA, but also more surprisingly with an increased release in the PFC which could explain their enhanced working memory ability. This apparent discrepancy between our electrophysiology data and the increased cortical release of dopamine is likely due to the fact that only a minority of the dopamine cells in the VTA project to the PFC (Bjorklund and Lindvall, 1984).

Dysregulation of the mesocorticolimbic system is a key feature of several stress-related behavioural psychopathologies, including addiction and depression that develop with a high interindividual variation that is not fully understood (Robinson and Berridge, 2000; Russo *et al*., 2012). The differences of cortical/subcortical dopaminergic balance between the higher and lower ranked individual we reported, may provide a physiological ground underlying differential vulnerability to psychopathology-like phenotypes. Indeed, the reduction in locomotor sensitization to cocaine in highest ranked individuals is coherent with the reduction of VTA bursting activity in these individuals (Runegaard *et al*., 2018). Repeated social defeat in mice has been intensively used as a preclinical model of depression. In a subset of animals, so-called susceptible, this chronic stress induces enduring anxiety and social avoidance that depends on enduring increase of VTA dopamine activity (Cao *et al*., 2010). Optogenetic stimulation of VTA neurons projecting to the NAcc induces a susceptible phenotype whereas optogenetic inhibition induces resilience (Chaudhury *et al*., 2013). We demonstrated that lowest ranked mice, with higher VTA dopamine tone are more likely to develop social aversion following ten days of repeated defeats. Two other studies made observations that differ from ours on the consequences of repeated social defeat depending on social rank. Lehmann *et al*. (2013) did not observe a correlation whereas Larrieu *et al*. (2017) observed the opposite (*i.e*. resilience for lower ranked individuals). Differences may reside in the intensity of the defeats to which individual were exposed, the lower number of R1 and R4 tested (8 here *vs* 4) and the fact that pooled data from ranks 1 and 2 were compared to that of ranks 3 and 4 in the Larrieu’s study. Furthermore, our experiment was performed on tetrads established for 5 months that have been tested regularly to ensure their stability over time. This repeated solicitation of animals in the context of chronic social competitions may have reinforced the phenotypes of each rank.

Studies in human and animals suggested the existence of a correlation between social rank and differences in stress hormones. Elevated circulating glucocorticoids are usually associated with subordinate status in non-mammals, and mammals including rodents and primates, although conflicting results have been reported (Sapolsky, 2004; Sapolsky, 2005; Creel *et al*., 2013; Cavigelli and Caruso, 2015). In human, the common perception that dominant individuals may have higher glucocorticoid levels has been challenged. Higher socio-economic status (SES) has been linked to lower evening glucocorticoid levels (Cohen *et al*., 2006). Studies in military leaders, as well as in influential individuals from a Bolivian forager-farmer population, different for individuals with higher SES, showed lower glucocorticoid levels (Sherman *et al*., 2012; von Rueden *et al*., 2014). We studied the social fate of mice deprived of the glucocorticoid receptor gene in dopamine innervated neurons. This targeted mutation clearly promotes higher social ranking in our tetrads. This result is consistent with our recent observation made on mice raised by two (Papilloud *et al*., 2020). Interestingly, we also showed that these mice exhibit a lower VTA dopamine cells bursting activity (Ambroggi *et al*., 2009), a decreased sensitization to cocaine (Barik *et al*., 2010) and a shift toward resiliency following repeated social defeat (Barik *et al*., 2013). These phenotypes are strikingly similar to that of R1 individuals suggesting that stress response and its impact on dopamine pathway might play a principle organizational role in shaping the behavioural trajectories leading to the establishment of social ranking. This hypothesis is reinforced by our data showing increased social ranking upon VTA dopamine neurons inhibition. In contrast, optogenetic stimulation of VTA dopamine neurons seems to favour dominant behaviour for competitive access to reward, which could rather reflect the role of this brain region in reward processing (Lozano-Montes *et al*., 2019). Importantly, in our experiment, whereas a transient reduction of dopamine activity during dominance measurement in tube-test has no detectable effect, a permanent reduction during the formation of the social organization upwards rank attainment, suggesting that this activity does not affect the expression of dominance but rather shapes its establishment. This process may occur under the continuous influence of the glucocorticoid stress-response.

## Material and methods

### Animals

C57BL/6JRj, 129/SvEv, and CD1 male mice of 6 weeks were purchased from Janvier (Le Genest-Saint-Isle, France) and housed under standard conditions, at 22°C, 55% to 65% humidity, with a 12-hour light/dark cycle (7 am/7 pm) and free access to water and a rodent diet. BAC-DATiCre*fto* mice (Turiault *et al*., 2007) were heterozygous and backcrossed on a C57BL/6J background. *Nr3c1* (*GR*) gene inactivation was selectively targeted in dopaminoceptive neurons (*Nr3c1*^loxP/loxP^;Tg:D1aCre (Lemberger *et al*., 2007), here after designed GR^D1aCre^), as described in Ambroggi *et al*. (2009). Experimental animals were obtained by mating Nr3c1^loxP/loxP^ females with Nr3c1^loxP/loxP^;Tg:D1aCre males. Half of the progeny were mutant animals the other half were control littermates. When required, thymus and adrenal glands of animals were dissected and weighted after fat tissue removal under a binocular loupe. Experiments were performed in accordance with the European Directive 2010/63/UE and the recommendation 2007/526/EC for care of laboratory animals and approved by the Sorbonne Université committee for animal care and use.

### Stereotaxic injections

Stereotaxic injections were performed using a stereotaxic frame (Kopf Instruments) under general anaesthesia with xylazine and ketamine (10 mg kg^−1^ and 150 mg kg^−1^, respectively). Anatomical coordinates and maps were adjusted from Watson and Paxinos, 2010. The injection rate was set at 100 nl min^−1^. To express hM4D in VTA dopamine neurons, we injected BAC-DATiCrefto mice (6 weeks old) with AAV8-hSyn-DIO-hM4D-mCherry (300 nl L^−1^, titration 10^12^ particles ml^−1^, University of North Carolina, USA vector core facility) bilaterally in the VTA (AP −3.2 mm, ML ±0.6 mm, DV −4.7mm from the bregma). For control, C57BL/6J mice issued from the same breedings than BAC-DATiCrefto mice, but not carrying the transgene, were injected similarly but with an AAV8-hSyn-GFP (titration 10^12^ particles ml^−1^, University of North Carolina, USA vector core facility). Animals were given a 4 weeks’ recovery period to allow sufficient viral expression.

### Constitution of tetrads

Mice were weighted upon arrival and were then grouped by four (tetrads) gathering mice of similar weights. When behavioural testing was performed before the constitution of the tetrads, mice were singly housed for one week. Mice were regularly weighted. For tetrads including GR^D1aCre^ mutant mice, animals were genotyped at 4 weeks of age. Tetrads were formed with animals unfamiliar to each other issued from different litters, grouping one mutant with three age-matched control mice (GR^loxP/loxP^).

### Social rank identification

#### Tube-test

Mice gathered by groups of four individuals for two to four weeks were first trained to move forward a transparent Plexiglas tube (diameter, 2.5 cm; length, 30 cm) for 2 consecutive days, performing 8 trials the first day and 4 the second one. Each individual alternatively entered the tube from right and left extremities and was let for a maximum of 30 seconds to exit the tube at the opposite end. After 30 seconds if still present within the tube, the mouse was gently pushed out. The diameter of the tube allowed passing one individual but and did not permit it to reverse direction. During the following days, social ranks were assessed daily through the six possible pairwise confrontations in the tube, performing for each a trial composed of 3 confrontations. Two mice were simultaneously introduced within the tube from the 2 opposite ends taking care that they met in the middle of the tube. The first mouse to exit the tube was designed as the loser of the contest. The individual that won at least 2 confrontations was ranked higher. Mice were classified from rank 1 (3 wins) to rank 4 (no win). Contests exceeding 2.5 minutes were stopped and immediately repeated. After each trial, the tube was cleaned with 20% ethanol and dried. Among 84 tetrads analysed, we always observed a non-ambiguous ranking. The order of confrontations was randomized day after day using a round-robin design. Social ranks were initially assessed during a minimum period of 6 days and considered stable if both ranks 1 and 4 were stable for the last three days. Tetrads that did not reach this criterion were analysed further, until reaching three days stability for ranks 1 and 4. All 84 tetrads reached stability within 12 days. Social rank was repeatedly analysed every three to four weeks for a minimum of three consecutive days.

For tetrads composed of one mouse stereotaxically injected with AAV8-hSyn-DIO-hM4D-mCherry and three mice with AAV8-hSyn-GFP, once both ranks 1 and 4 were stable for three consecutive days, tube test was pursued for five more days but each individual received an injection of CNO (*i.p*. 1mg kg^−1^), 30 minutes before tube-test. Three weeks later, tetrads were treated for 24h with CNO in drinking water (10 mg L^−1^), then dismantled and new tetrads were formed with similar composition but with individuals unknown to each other and were permanently treated with CNO in drinking water (10 mg L^−1^). After two weeks, social ranking was determined by tube test.

#### Territory urine marking assay

R1 and R4 mice from a tetrad were placed in an empty PVC box (42×42×15 cm), separated by a central transparent perforated Plexiglas divider and were let free to explore and mark their own territory for 2 hrs. Absorbent paper (Whatmann), partially covered by fresh sawdust was set in the bottom of each compartment to collect urine deposited by mice during the session. Absorbent paper was then pictured under UV light (312 nm). Both the number of urine marks and the total area of urine marks were quantified.

#### Warm spot assay

Tetrads were placed in a transparent plastic cage (35×20×18 cm) without litter, placed on ice (bottom cage temperature 4 °C). 20 minutes later, a warm plate (11×9 cm, 28-30 °C) was introduced on the floor of the cage, at a corner. Mice activity was recorded for 20 min, and warm plate occupancy, by each individual, scored by an experimenter blind to conditions.

### Spontaneous locomotor activity in open field

Mice were placed in a corner of a squared PVC white box (42×42 cm, 15 cm depth), and let free to explore for 10 min, under 50 lux. A video camera system placed above enabled the automatic quantification of locomotor activity (Noldus Ethovision 11.0 XT).

### Anxiety-like behaviour

#### Dark-Light box

The dark-light box apparatus consisted of a plastic rectangular box (45×20 cm, 25 cm high) divided into a white compartment (30 cm, open) and a black compartment (15 cm, covered with a removable lid), that communicate through a central door (5×5 cm). Animals were initially placed into the black compartment, and exploration recorded for 10 min, under 30 lux. The time spent in each compartment was blindly scored by two experimenters.

#### Elevated O-maze

The maze consisted of a circular path (width 5.5 cm, outer diameter 56 cm) elevated 30 cm above the floor and made of black PVC. It was divided in four sections of equal lengths, two opposite bordered with bilateral black plastic walls (15.5 cm high) and two open ones. Mice were positioned at one extremity of a closed section, the head directed inward, under 50 lux in the open sections and 10 lux in the closed one. Their exploration was recorded for 10 minutes and the time spent into closed and open sections was blindly scored by two experimenters. A mouse was considered to be in a section when the 4 paws were introduced.

### Despair, forced swim test

Glass cylinders (40 cm tall, 12 cm diameter) were filled with tepid water (23°C) until reaching a depth of 10 cm. Mice, placed on a large spoon, were gently introduced into cylinders and videotaped for 6 min. Cumulative length of time of immobility, balance and escape movements were blindly scored. Escape behaviour was defined as movements involving the 4 paws of the animal beating against the wall of the cylinder mimicking a climbing-like behaviour. Balance movements refer to brief movements involving mainly only the 2 posterior paws of the animal and aiming to displace in water without trying to climb up the cylinder’s wall. Mice were considered immobile when floating passively, doing neither escape nor balance movements. The experiment was repeated 24 h later when planed in the experimental design.

### Sociability, three-chambers test

Sociability was measured under 50 lux in a rectangular box containing three chambers (30×20, 15 cm high for each compartment) with removable doors (5×5 cm) at the centre of each partition. In the opposite sides of the 2 lateral compartments, 2 clear perforated plastic boxes (10×7 cm, 7 cm high) were placed. One contained an unfamiliar adult male mouse (C57BL/6J), the other was left empty. During habituation phase (5 min), the challenged individual was placed in closed central compartment. Doors were then opened and the mouse free to explore the display for 5 min. The sessions were recorded, and the close interaction time with the empty box and with the box containing an unknown congener were blindly scored. The interaction time was defined as the periods during which the animal was oriented with the head towards the box, and in direct contact with it. To measure the preference for social novelty, and social memory, the mouse was let, closed, in the chamber containing the social cue for 5 min. It was then placed again in the central chamber, free to investigate the three compartments. The time length spent in close interaction with boxes containing, either a familiar mouse, previously encountered, or an unfamiliar one was scored, for 5 minutes session.

### Aggressiveness, resident intruder challenge

Ranks 1 and 4 males were individually housed for 48 hours before starting the resident intruder test. 10 weeks old 129/SvEv males (Janvier laboratory, Saint-Berthevin, France) were used as intruders. The experimental sessions were carried out between 9.30 am and 3 pm. The intruder was placed in the cage of the challenged mouse and social interactions were videotaped for 20 minutes for ulterior manual scoring by an experimenter blind to the conditions. A second session was repeated 24 h later with a new intruder. Sessions in which either R1 or R4 individuals from on tetrad displayed aggressive behaviour were scored.

### Non-matching to sample T-maze task

The test was performed as previously described (Sigurdsson *et al*., 2010). Briefly, mice underwent for 3 days a moderate food reduction (2 g/mouse/day), taking care not to go below 85% of their initial weight. Animals were then trained on a spatial working memory task (non-match-to-sample task) in a T-maze (61 cm large x 51 cm width x 15 cm high, with a path 11 cm large). Mice were habituated to the maze for two days during which they had 15 minutes to collect food pellets (20 mg dustless sugar pellets, Bioserv). The next three days, mice had to complete 4 forced runs each day, during which one of the two arms were alternatively closed in order to habituate to the guillotine doors (Fig. 2D). Mice were then daily tested on 10 trials per day. Each trial consisting of two runs, a forced run and a choice run. At the beginning of the trial both arms are baited. In the forced run the right or left arm is randomly chosen to be opened, while the other arm is closed. At the beginning of the forced run, the mouse was placed at end of the longest T-maze arm. After running down this arm, it could enter into the open goal arm and have access to a food reward. Once the mouse reached back the starting arm, it was blocked by a door at its end for a delay of 6 s. Then started the choice run. During it, the mouse ran down the centre arm, where it had to choose between the two open goal arms. To obtain a reward, animals were required to enter the non-visited arm during the sample phase. This was scored as a correct choice. Animals were exposed to daily sessions of 10 trials, until they reached a criterion performance, defined as having a minimum of seven correct choices a day, for three consecutive days. The inter-trial time was 45 s.

### Social defeat and interaction paradigms

Social defeat was performed as previously described (Barik *et al*., 2013). Six months old CD1 breeder male mice were screened for their aggressiveness. 6 months old individuals were subjected to 10 consecutive days of social defeat with new encounters. Each defeat consisted of 5 minutes physical interactions with a resident CD1 mouse, followed by a 24 h exposure to the CD1 in its home cage but separated by a perforated transparent plastic wall which allowed visual, auditory, and olfactory communication whilst preventing physical contact. Social interaction was first performed the day before the first social defeat (pre-defeat) and performed again 24 h after the last social defeat (post-defeat). Challenged mice were placed for 150 seconds in a plastic white open-field (42×42 cm, 30 cm high, 20 lux) containing an empty transparent and perforated plastic box. Mice were rapidly removed and an unfamiliar CD1 mouse was placed in the box, and the challenged mouse re-exposed to the open field for 150 seconds. Sessions were recorded and the time spent in direct interaction with the boxes were manually quantified by an experimenter blind to conditions.

### Locomotor sensitization to cocaine

Locomotor sensitization to cocaine was conducted on 3 months old R1 and R4 individuals. Mice were placed in a circular corridor (4.5-cm width, 17 cm-external diameter, 30-50 lux) crossed by four infrared captors (1.5 cm above the base), equally spaced (Imetronic, Pessac, France). The locomotor activity was automatically quantified by counting the quarters of turn travelled by the mouse that corresponded to the interruption of two successive beams. Animals were habituated to the apparatus for 3 hours during 3 consecutive days and received a saline injection on days 2 and 3 (NaCl 0.9% saline solution, 10 ml kg^−1^, *i.p.*). On the five following days, mice were placed in the apparatus for 90 min, then injected with cocaine hydrochloride (Sigma-Aldrich, 10mg kg^−1^ *i.p*.) and left inside 180 minutes after injection. Following 7 days of withdrawal, mice received a challenge injection of cocaine (10mg kg^−1^ *i.p*). At the end of each session, mice were placed back in their tetrads. Social ranks were tested at the end of the experiment, and only mice of R1 and R4 that did not change were considered for the analysis. The behavioural sensitization experiment has been carried out from 9 am to 13 pm.

### In vivo electrophysiological recordings

3 to 5 months old R1 and R4 mice were anesthetized with chloral hydrate (8%), 400 mg kg^−1^ *i.p*. supplemented as required to maintain optimal anaesthesia throughout the experiment and positioned in a stereotaxic frame. A hole was drilled in the skull above midbrain dopaminergic nuclei (coordinates: 3.0 ± 1.5 mm posterior to bregma, 1 ± 1 mm [VTA] lateral to the midline, Watson and Paxinos, 2010). Recording electrodes were pulled from borosilicate glass capillaries (with outer and inner diameters of 1.50 and 1.17 mm, respectively) with a Narishige electrode puller. The tips were broken under microscope control and filled with 0.5% sodium acetate. Electrodes had tip diameters of 1-2 μm and impedances of 20–50 MΩ. A reference electrode was placed in the subcutaneous tissue. The recording electrodes were lowered vertically through the hole with a micro drive. Electrical signals were amplified by a high-impedance amplifier and monitored with an oscilloscope and an audio monitor. The unit activity was digitized at 25 kHz and stored in Spike2 program. The electrophysiological characteristics of dopamine neurons were analysed in the active cells encountered when systematically passing the microelectrode in a stereotaxically defined block of brain tissue including the VTA (1). Its margins ranged from −2.9 to −3.5 mm posterior to bregma (AP), 0.3 to 0.6 mm (ML) and −3.9 to −5 mm ventral (DV) (Grace and Bunney, 1984). Sampling was initiated on the right side and then on the left side. Extracellular identification of dopamine neurons was based on their location as well as on the set of unique electrophysiological properties that distinguish dopamine from non-dopamine neurons *in vivo*: (i) a typical triphasic action potential with a marked negative deflection; (ii) a long duration (>2.0 ms); (iii) an action potential width from start to negative trough >1.1 ms; (iv) a slow firing rate (<10 Hz and >1 Hz). Electrophysiological recordings were analysed using the R software (^49^). Dopamine cell firing was analysed with respect to the average firing rate and the percentage of spikes within bursts (%SWB, number of spikes within burst divided by total number of spikes). Bursts were identified as discrete events consisting of a sequence of spikes such that: their onset is defined by two consecutive spikes within an interval <80 ms whenever and they terminate with an inter-spike interval >160 ms. Firing rate and % of spikes within bursts were measured on successive windows of 60 s, with a 45 seconds overlapping period. Responses to nicotine are presented as the mean of percentage of firing frequency variation from the baseline ± SEM. For statistical analysis, maximum of firing variation induced by nicotine occurring 180 seconds after the injection are compared to spontaneous variation from the baseline occurring 180 seconds just before the injection by non-parametric Mann-Whitney test.

### Quantification of dopamine and DOPAC

Animals were decapitated and brains were rapidly dissected and frozen at −12°C on the stage of a Leitz-Wetzlar microtome. Coronal sections (300 μm thick) were cut and placed onto the refrigerated stage. Three dopaminergic terminal fields were assayed: the mPFC, the CPu and the NAcc. For each structure, two or four tissue punches (1 mm diameter) from two consecutive sections were taken bilaterally and each side analysed separately for the CPu, the NAcc and mPFC, respectively. Tissue punches were immersed into 50 μl of 0.1 N HClO_4_ containing Na_2_S_2_O_5_ (0.5%), disrupted by sonication and centrifuged at 15000 g for 20 min. Aliquots (10μl) of supernatant were diluted with high pressure liquid chromatography mobile phase and injected into a reverse-phase system consisting of a C18 column (HR-80 Catecholamine 80 x 4.6 mm, Thermo Scientific, USA) and a 0.1 M NaH_2_PO_4_ mobile phase containing 1-octanesulfonic acid (2.75 mM), triethylamine (0.25 mM), EDTA (0.1 mM), methanol (6 %) and adjusted to pH 2.9 with phosphoric acid. Flow rate was set at 0.6 mL/min by an ESA-580 pump. Electrochemical detection was performed with an ESA coulometric detector (Coulochem II 5100A, with a 5014B analytical cell; Eurosep, Cergy, France). The conditioning electrode was set at – 0.175 mV and the detecting electrode at + 0.175 mV, allowing a good signal-to-noise ratio. External standards were regularly injected to determine the elution times (9.8 and 21.5 min) and the sensitivity (0.3 and 0.4 pg), for DOPAC and dopamine respectively.

### Immunostaining

Following CNO treatment, anesthetized mice were perfused (intra cardiac) with 4% paraformaldehyde. Brains were removed, post-fixed over-night in the same solution and sliced with a vibratome (30 μm). Sections containing SN and VTA were incubated with antibodies directed against tyrosine hydroxylase (mouse monoclonal, 1:1000, Merck-Millipore MAB318) and mCherry (rabbit polyclonal, 1:500, Abcam ab167453) overnight at 4°C with constant shaking. Sections were then incubated for 2 h at room temperature with anti-mouse Alexa 488 (Invitrogen A-1101, 1:500) and anti-rabbit Cy3 (Invitrogen A-10520, 1:500) antibodies. Sections were mounted using Vectashield with DAPI (Vertorlabs) and analysed by fluorescence microscopy.

### Statistical Analysis

Data are expressed as mean ± SEM. Statistical analyses were performed by using Mann-Whitney non-parametric test and two-way ANOVA. When primary effect was found to be significant, post hoc comparisons were made using a Bonferroni/Dunn test. Differences of P ≤0.05 were considered statistically significant. Statistical analyses were carried out using PRISM software.

## Data and materials availability

The that support the findings in this study are available from the corresponding authors upon request. Source data are provided with this paper.

## Acknowledgements

This project has been supported by grants from the LabEx BioPsy, the French Agence Nationale de la Recherche (ANR3053NEUR31438301 to FT and ANR-14-CE35-0029-01 to SP), by the Foundation for Medical Research (FRM Equipe grant DEQ20140329552 to FT and DEQ20180339159 to JB), by the INCA (TABAC grant to PF, JB and FT), Université Côte d’Azur and Sorbonne Université. The authors warmly thank S. Bhattacharya, C Sandi, L. Amar and S. Vyas for helpful discussions and S.B., L. A and S.V. for critical reading of the manuscript. We also thank F. Machulka and the IBPS mouse facility for their technical assistance.

## Contributions

F.T. and S.P conceived and co-supervised the study. F.T., S.P, D.B. and P.F. designed the experiments, acquired and analysed the data. D.B., C.V., C.N., S.P., A.Z., A.C.M., S.M., A.F., J.P.T., T.C., J.B. and F.M. acquired and analysed the data. F.T., D.B. and S.P, wrote the manuscript.

---

**Extended data Figure 1.**
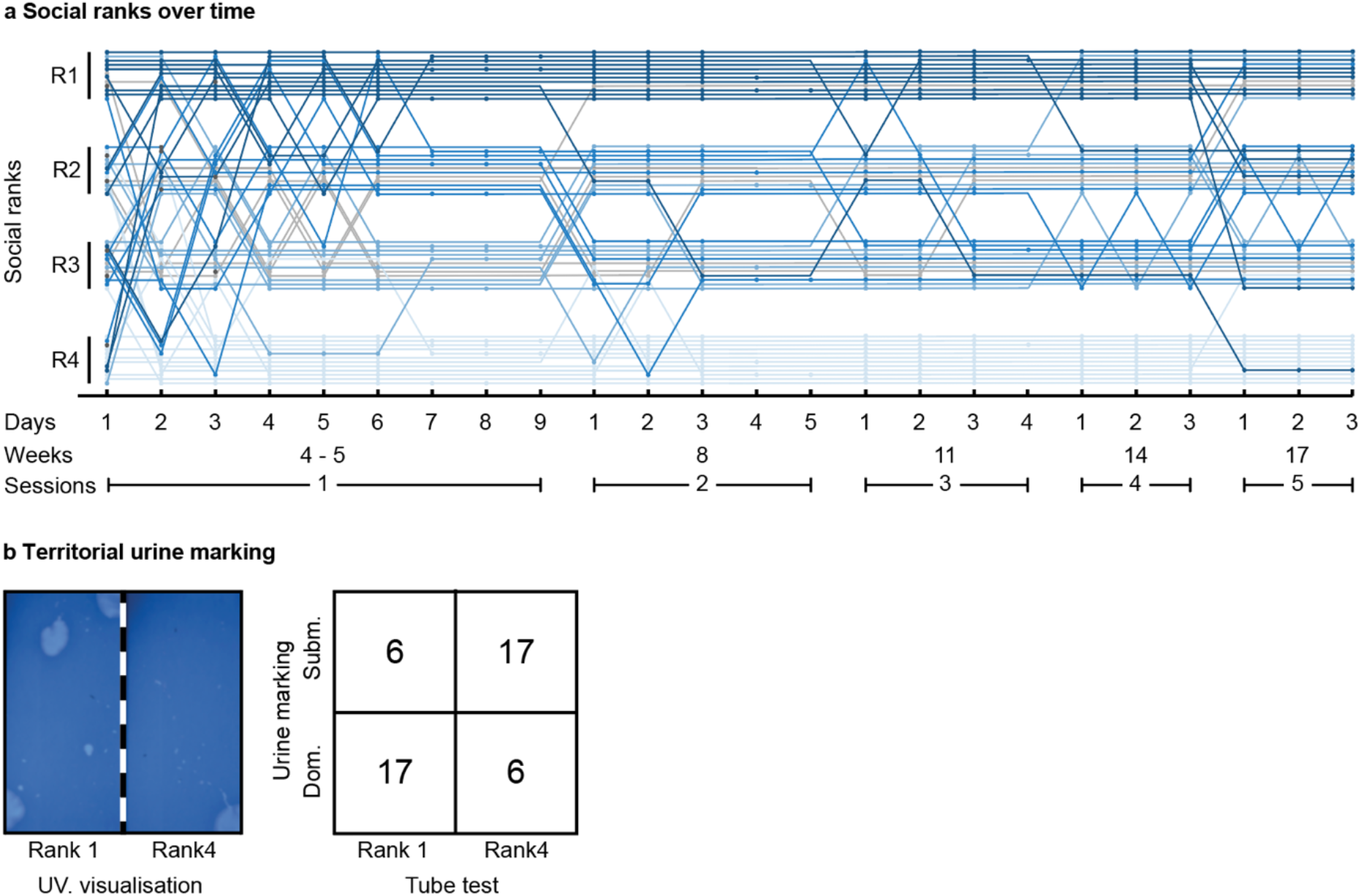
Stability of hierarchical classification and territoriality. **a**, Rank assessment in tube-test is pictured with daily resolution for 12 tetrads. As in Fig. 1, each line corresponds to an individual mouse and indicates the tetrad to which it belongs to (1 to 12). The blue intensity indicates it rank attainment after the first session. R1-4 indicates its social rank. Dots on the lines indicate that a tube-test was performed during the corresponding day. Note that there is no dot after hierarchy has reached the stability criterion during a session (*i.e*. that R1 and R4 mice were stable for three consecutive days, with an initial period of at least 6 days of testing). Grey lines correspond to mice of intermediate ranks that did not reach stability at the end of the first session. On days 1, 2 and 3, dark grey dots indicate individuals for which the rank could not be determined. **b**, Territorial urine marking reflects ranking obtained in tube-test. Left : representative picture of a urine marking during a 2 h confrontation between a R1 and a R4 mice, visualized under UV light. Right : contingency table of the ranking correspondences between the tube- and the urine marking-tests. Dom. : dominant; Subm : submissive. Fisher’s exact test, two-tailed, **p<0.01, n=23 per group.

**Extended data Figure 2.**
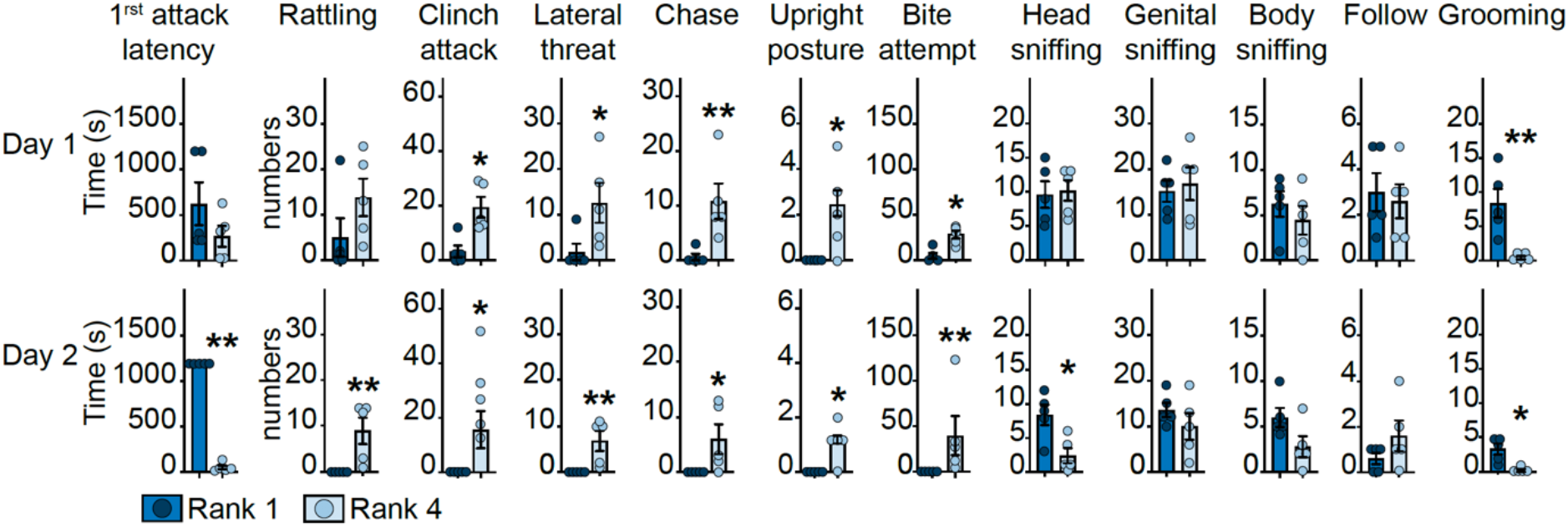
Aggressiveness of rank 1 and rank 4 mice. The latencies to the first attack during resident-intruder challenges were scored for rank 1 (dark blue) and rank 4 individuals (light blue, left graph). The occurrences of aggressive and not aggressive behaviours were scored during 20 min. Data are expressed as mean ± SEM. Statistical analyses were performed by using Mann-Whitney non-parametric test. *p<0.05; **p<0.01 (n=5 per group).

**Extended data Figure 5.**
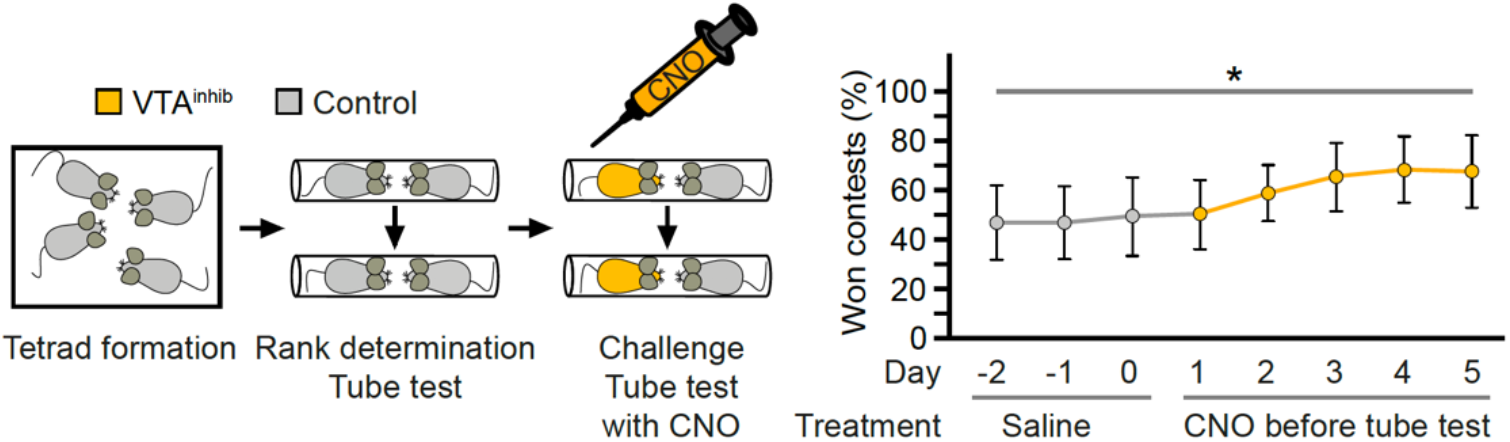
Reducing VTA dopamine neurons activity during tube test only does not affects social ranking. Left, tetrads were constituted with one mouse expressing CNO-dependent hM4D and (VTA^inhib^) and three expressing GFP in the VTA (control) mice, in absence of CNO treatment. After two weeks, social hierarchy was determined (n=7 cages). During the five following days, tube tests were repeated with mice receiving an injection of CNO, 30 minutes before. The right graph pictures the percentage of won tube test contests by VTA^inhib^ mice on the last three days without CNO treatment (days −2 till day 0, saline) and the five following days, after a CNO injection. Data are expressed as mean ± SEM. Repeated ANOVA. * p<0.05.

